# Disulfide Crosslink Density is a Key Determinant in the Formation, Dynamics, and Maturation of Cysteine-rich Designer Biomolecular Condensates

**DOI:** 10.64898/2026.11.03.763743

**Authors:** Malay Mondal, Windfield S. Swetman, Conner T. Allen, Harrison Livingston, Tristan D. Clemons, Vijayaraghavan Rangachari

**Affiliations:** Department of Chemistry and Biochemistry, School of Mathematics and Natural Sciences, University of Southern Mississippi, Hattiesburg, MS, USA; School of Polymer Science and Engineering, University of Southern Mississippi, Hattiesburg, MS, USA

**Keywords:** Biomolecular condensates, Liquid-Liquid phase separation, membraneless organelles, redox, cysteines, disulfide bonds, viscoelasticity

## Abstract

Biomolecular condensates form through weak, multivalent, non-covalent interactions, but the role of covalent crosslinks in shaping their assembly and material properties remains poorly understood. We previously showed that disulfide bonds between cysteines in designer phase-separating peptides (PSPs) act as redox-reversible crosslinks that lower the saturation concentration (*C*_sat_) and help maintain condensate fluidity. Here, we ask which sequence features govern this behavior. We built two peptide libraries: one varying the inter-cysteine spacer length (2–10 residues) and one varying cysteine density (0–33%). Inter-cysteine spacers tuned condensate properties only modestly. Longer inter-cysteine lengths raised *C*_sat_ from 300 to 900 μM, showed greater fluidity (as assessed by fluorescence recovery after photobleaching (FRAP) recovery rate, and resisted loss of fluidity during maturation; all inter-cysteine length variants dissolved and reformed reversibly over three redox cycles. Cysteine density, in contrast, was the dominant control parameter. Increasing cysteine density lowered *C*_sat_ about 60-fold (1.5 mM to 25 μM) and reduced the fluidity by an order of magnitude. Its effect on fluidity was non-monotonic: condensates with up to ∼14% cysteine density stayed liquid-like for 10 days, whereas those with 21–33% cysteine density in the peptide showed slower coalescence. Condensates with the highest densities also resisted dissolution under reducing conditions and showed sharply reduced uptake of a protein cargo (eGFP; ∼60% to ∼9%) and a small molecule (TMR, ∼70% to ∼45%), consistent with smaller mesh size. Together, these results identify disulfide crosslink density as a key lever for programming condensate assembly, viscoelasticity, redox responsiveness, and porosity. The results offer key design principles both for understanding cysteine-rich disordered proteins in biomolecular condensates and for engineering redox-responsive biomaterials.

**GRAPHICAL ABSTRACT:** 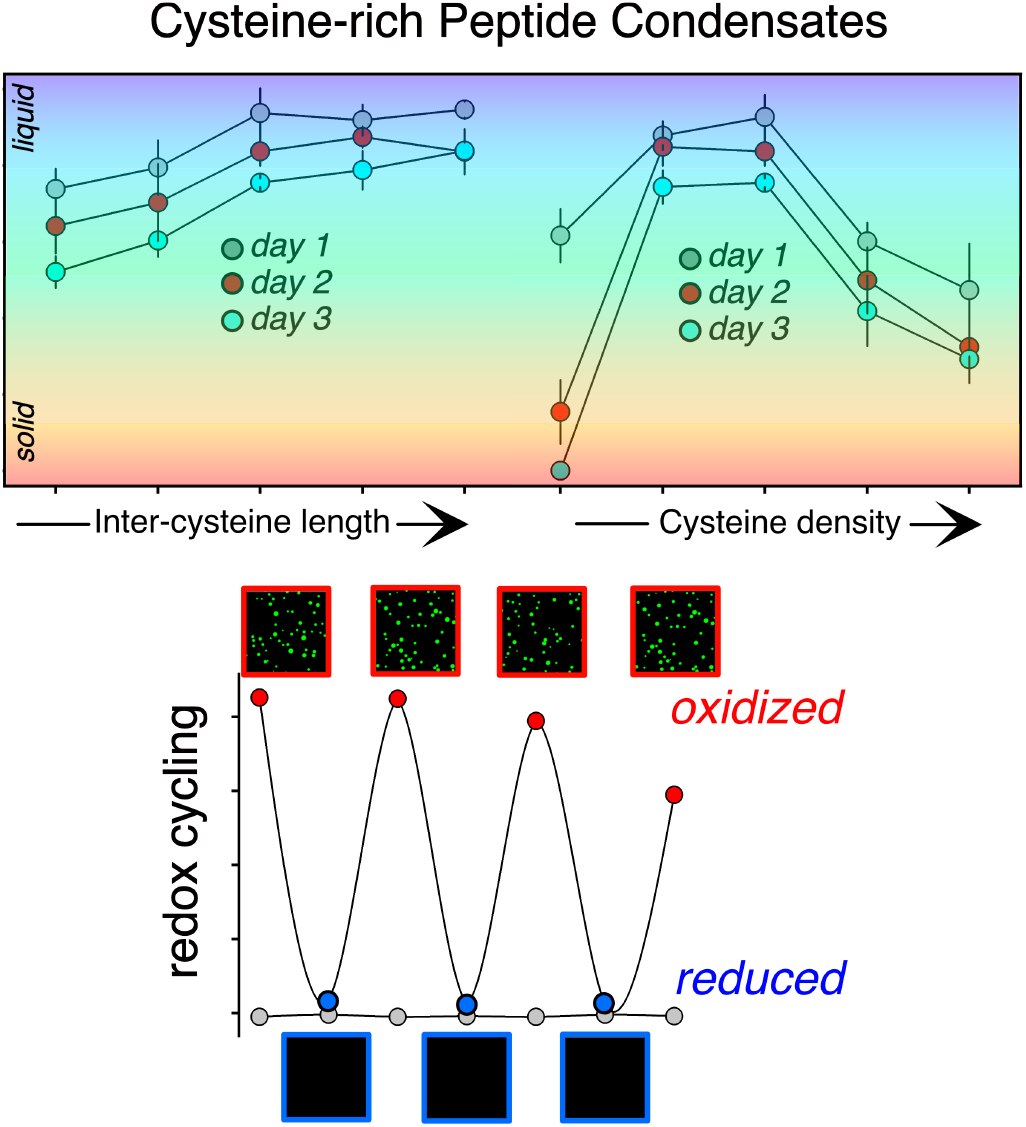

## INTRODUCTION

Biomolecular condensates (BCs), membraneless organelles predominantly formed by liquid–liquid phase separation (LLPS), are dynamic viscoelastic hubs enriched in proteins and nucleic acids.^1–8^ BCs participate in transient spatiotemporal reorganization, sequestering biomolecules into dense hubs to regulate a wide range of cellular functions.^5,9–13^ Needless to say, these viscoelastic condensates have garnered significant attention not only for their biological functions but also for their potential as biomaterials in many biotechnological applications.^7,14,15^ Investigations into the molecular determinants of BC development have revealed weak multivalent non-covalent interactions such as π–π, cation–π, or electrostatic interactions interspersed with disorder-promoting sequences drive BC formation.^13,16–20^

To understand the effect of long-range covalent bonds, if any, in condensate formation, we recently studied designer peptides with classical phase-separating sequences interspersed with cysteines.^21^ We found, that under oxidizing conditions, cysteines not only promote BC formation but also sustain condensate viscosity by inhibiting percolation networks. These observations suggest that cysteine disulfide bonds serve as critical covalent cross-links, lowering the critical saturation concentration (*C_sat_*) and promoting condensate stability. Furthermore, under oxidizing conditions, disulfide bonds play a critical role in condensate formation; under reducing conditions, their abrogation leads to complete condensate dissolution, and this formation and dissolution are fully reversible under redox flux.^21^ We also showed that disulfide-crosslinked condensates can serve as effective cell-delivery vehicles for molecular cargo.^22^ Due to the reducing environment of the cellular cytoplasm, cargo-laden condensates dissolve upon entering cells, releasing cargo into the cytoplasmic milieu.

Our prior work on cysteine-rich phase-separating peptides (PSPs) established the significance of redox-reversible cysteine disulfide bonds in phase separation. In this study, we test the hypothesis that sequence properties such as the inter-cysteine distance (spacer length) and disulfide cross-link density (fraction of cysteines) affect condensate properties and dynamics. Our results show that cysteine density primarily influences *C_sat,_* driving robust condensate formation and fluid properties, while inter-cysteine lengths and arginine-tyrosine interactions contribute secondarily. Furthermore, cysteine density and condensate dynamics show a non-monotonic relationship, spanning Newtonian liquids to possibly viscoelastic gels. These findings identify the sequence features that govern condensates formed by cysteine-rich disordered proteins and establish design principles for engineering redox-responsive biomaterials.

## RESULTS

### Peptide Design

To systematically investigate how sequence features, specifically spacer length and cysteine density, influence phase separation, we designed two libraries of phase-separating peptides (PSPs) as shown in Figure 1. The first library, spacer-length variants, was designed by incrementally increasing the spacer residues, serine (S) and glycine (G), from 2 to 10 amino acids while keeping cysteine (C) and the weakly interacting nodes, tyrosine (Y) and arginine (R), constant (Figure 1a). The notations follow, with PSP-2 as our base peptide and (+) or (−) denoting spacer length increases or decreases {PSP-2(−2), PSP-2(−1), PSP-2, PSP-2(+1), and PSP-2(+2) in Figure 1a}. Serine and glycine residues predominate in disordered protein regions and thus serve as ideal spacer residues.^16,23^ Such a design would allow us to investigate interactions between arginine and tyrosine (i.e. cation–π interactions) and disulfide bond formation as a function of inter-cysteine lengths and the length between the interacting nodes. The second group, cysteine variants, varied cysteine density by increasing the number of cysteines in the sequence while keeping the inter-cysteine lengths and interacting nodes constant (Figure 1b). Again, we used PSP-2, consisting of two cysteines, as a base and varied the cysteine content from 0 to 5 {PSP-2(C0), PSP-2(C1), PSP-2(C2), PSP-2(C3), PSP-2(C4), and PSP-2(C5)}, reflecting the cysteine densities of 0 to 35% (Figure 1b). In these peptides, we systematically introduced cysteine residues at the N- and C-termini or within the central spacer region to assess how disulfide crosslink density contributes to condensate formation and its fluid and material properties. We synthesized all peptides as C-terminal amides and verified them by electrospray ionization mass spectrometry (Figure S1). Together, these two peptide libraries provide a minimal yet informative design space for dissecting the roles of spacer length and cysteine composition in driving phase separation.

**Figure 1.**
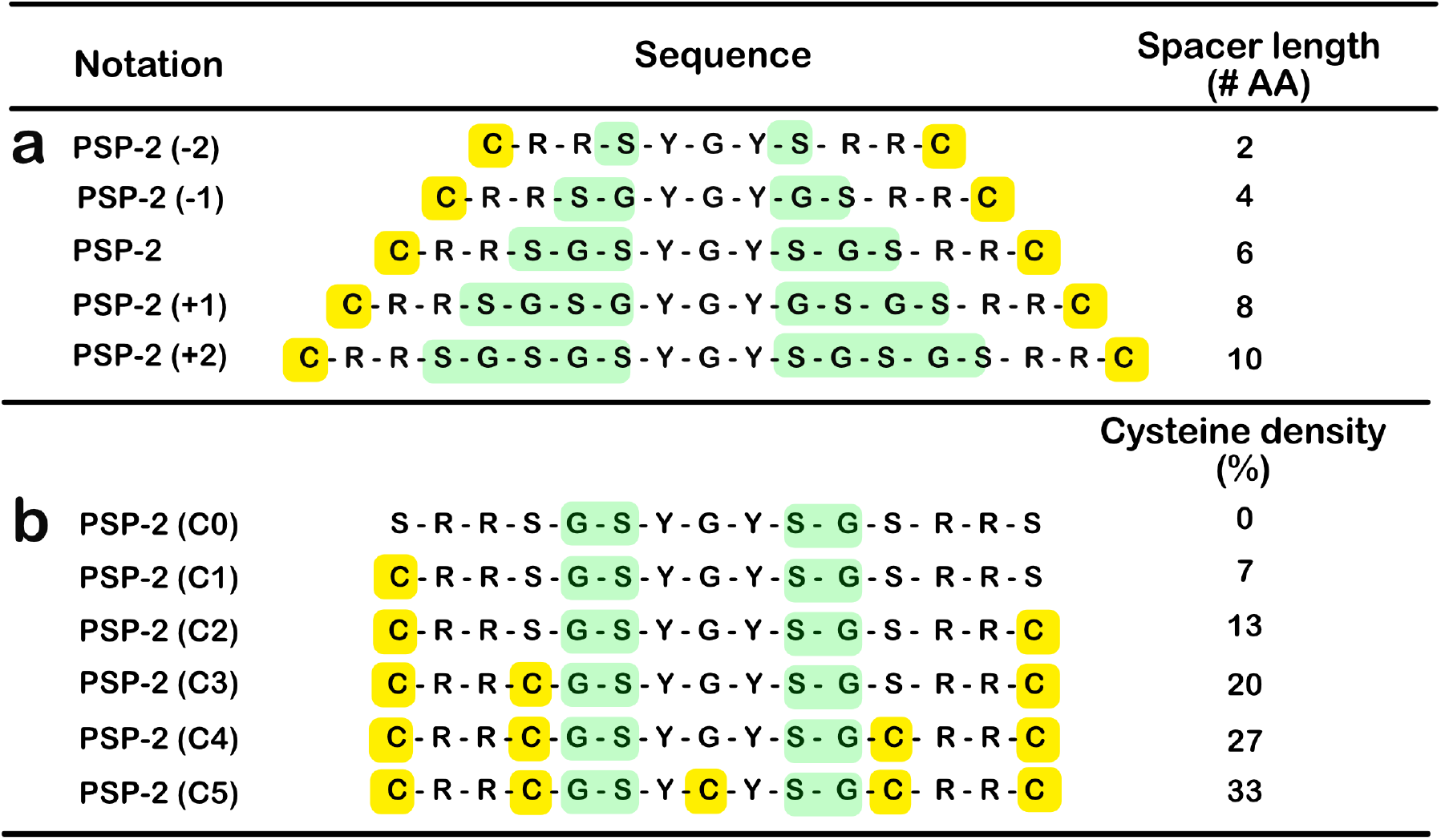
Phase-separating peptides used in this study. (a) Spacer-length variants designed by varying the number of glycine (G) and serine (S) spacer residues between constant arginine (R), tyrosine (Y), and cysteine (C) residues. The inter-cysteine spacer length is indicated as the number of amino acids. (b) Cysteine variants are designed by systematically interspersing cysteine residues within the peptide sequence, thereby varying the cysteine density from 0 to 33%.

### Cysteine Density Decreases Saturation Concentrations of Condensate*s*

To assess the effects of inter-cysteine spacer lengths and cysteine density on phase-separation behavior, we first quantitatively measured the saturation concentration (*C_sat_*) of PSP variants using sedimentation assays, as described in the Methods section. Briefly, we defined *C_sat_* as the point where theoretical and measured supernatant concentrations diverge in phase-separating (red circles) and non-phase-separating conditions (grey circles). Both peptide groups showed a monotonic increase in *C_sat_* with spacer length or cysteine density (Figure 2). While the *C_sat_* values for the spacer variants range from 300 μM for PSP-2(−2) to 900 μM for PSP-2(+2) (Figure 2a-e), they range between 25 μM and 1.5 mM for PSP-2(C5) to PSP-2(C1) (Figure 2f-j). As shown in Figure 2a–e, peptide concentrations measured under non-phase-separating conditions followed a mostly linear relationship with theoretical concentrations. In contrast, under phase-separating conditions, measured concentrations deviated from the control trend once a threshold concentration was reached, indicating condensate formation. The plot of *C_sat_* values against spacer length or cysteine density shows a monotonic relationship as expected (Figure 2k and 2l, respectively). Among the spacer variants, the shortest spacer, PSP-2(−2), showed the lowest *C_sat_* value (300 μM), while the longest spacer, PSP-2(+2), showed the highest *C_sat_* value (900 μM), with PSP-2(+1) as the only outlier from the linear relationship. This suggests that intramolecular sticker interactions and disulfide bond formation may dominate short inter-cysteine lengths, whereas intermolecular interactions may dominate peptides with longer inter-cysteine lengths. This hypothesis is also supported by results on cysteine density variants, which showed an inverse monotonic relationship between cysteine density and peptides with low or high *C_sat_* values, respectively (Figure 2l). However, unlike the spacer variants, the linear relationship between *C_sat_* and cysteine density had a lower slope, with the *C_sat_* range spanning more than two orders of magnitude, suggesting that disulfide crosslink density influenced phase separation more than spacer length.

**Figure 2.**
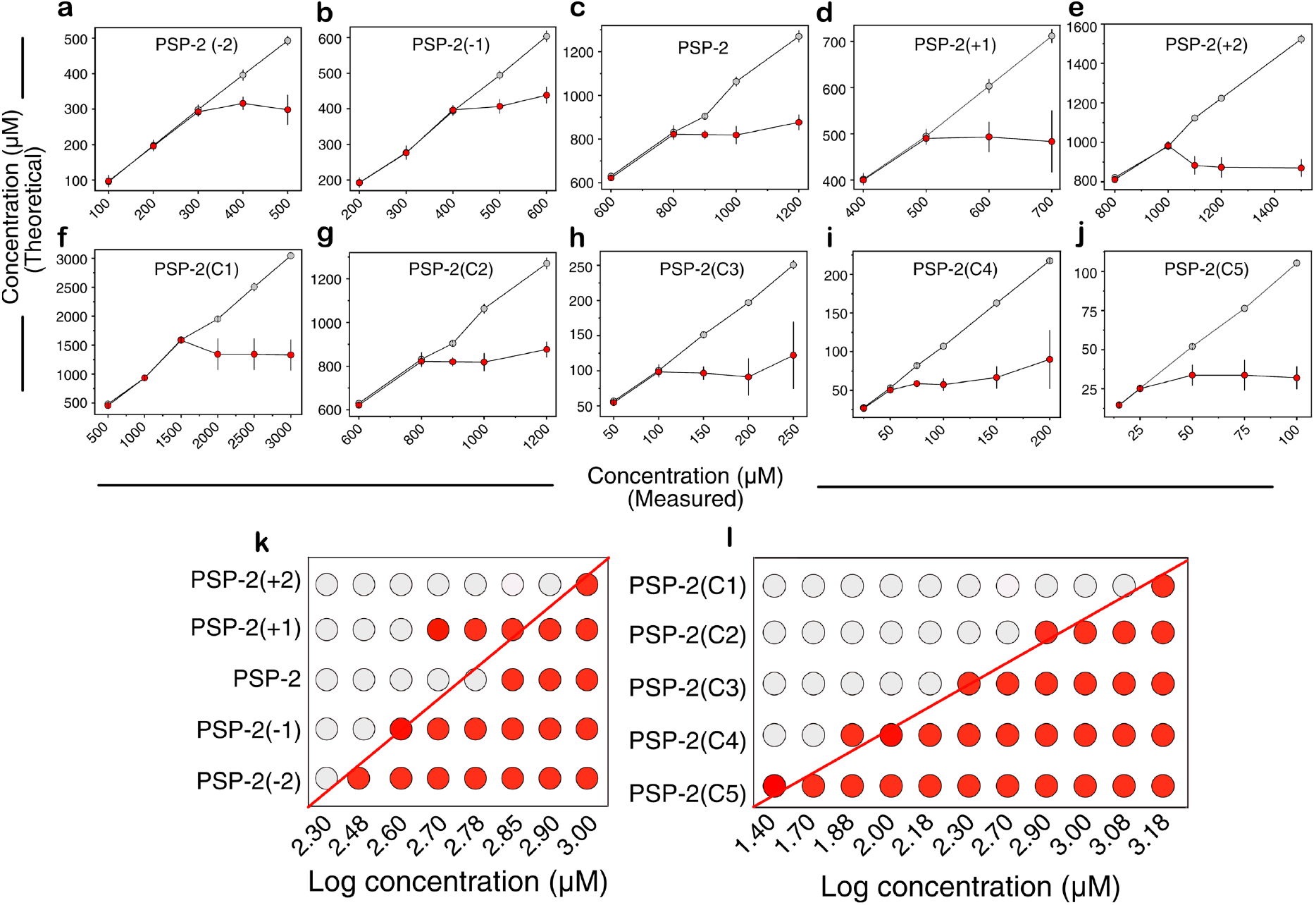
Phase boundaries of condensates formed by spacer-length and cysteine density variants. Saturation concentrations (*Csat*) were determined by sedimentation assay for PSP variants differing in spacer lengths and cysteine densities. (a–e): Measured peptide concentration in the dilute phase as a function of total peptide concentration for the spacer-length variants, PSP-2(−2), PSP-2(−1), PSP-2, PSP-2(+1), and PSP-2(+2). (f–j): Measured peptide concentration in the dilute phase as a function of total peptide concentration for the cysteine variants PSP-2(C1), PSP-2(C2), PSP-2 (C3), PSP-2(C4), and PSP-2(C5). Open gray circles represent concentration measurements under non-phase-separating conditions, while red circles represent measurements under phase-separating conditions. The inflection point at which phase-separating and non-phase-separating concentrations diverge was considered the *Csat*. (f and J): Derived *Csat* values vs. concentration plots for spacer-length and cysteine density variants, respectively. The red line shows the monotonic trend. Data displayed as mean ± standard deviation (n = 3).

Peptides differing in spacer length and cysteine density did not differ in droplet morphology. Both brightfield and confocal fluorescence images revealed well-defined, spherical droplets for both spacer-length and cysteine-density variants (Figure 3a-e and 3k-o, respectively). However, the two peptide groups differed in the droplet size formed immediately after incubation. While the spacer variants largely formed monodisperse droplets centered around 4 μm in diameter, the cysteine variants showed a wide size range from 1 to 6 μm (Figure 3f-j and 3p-t).

**Figure 3.**
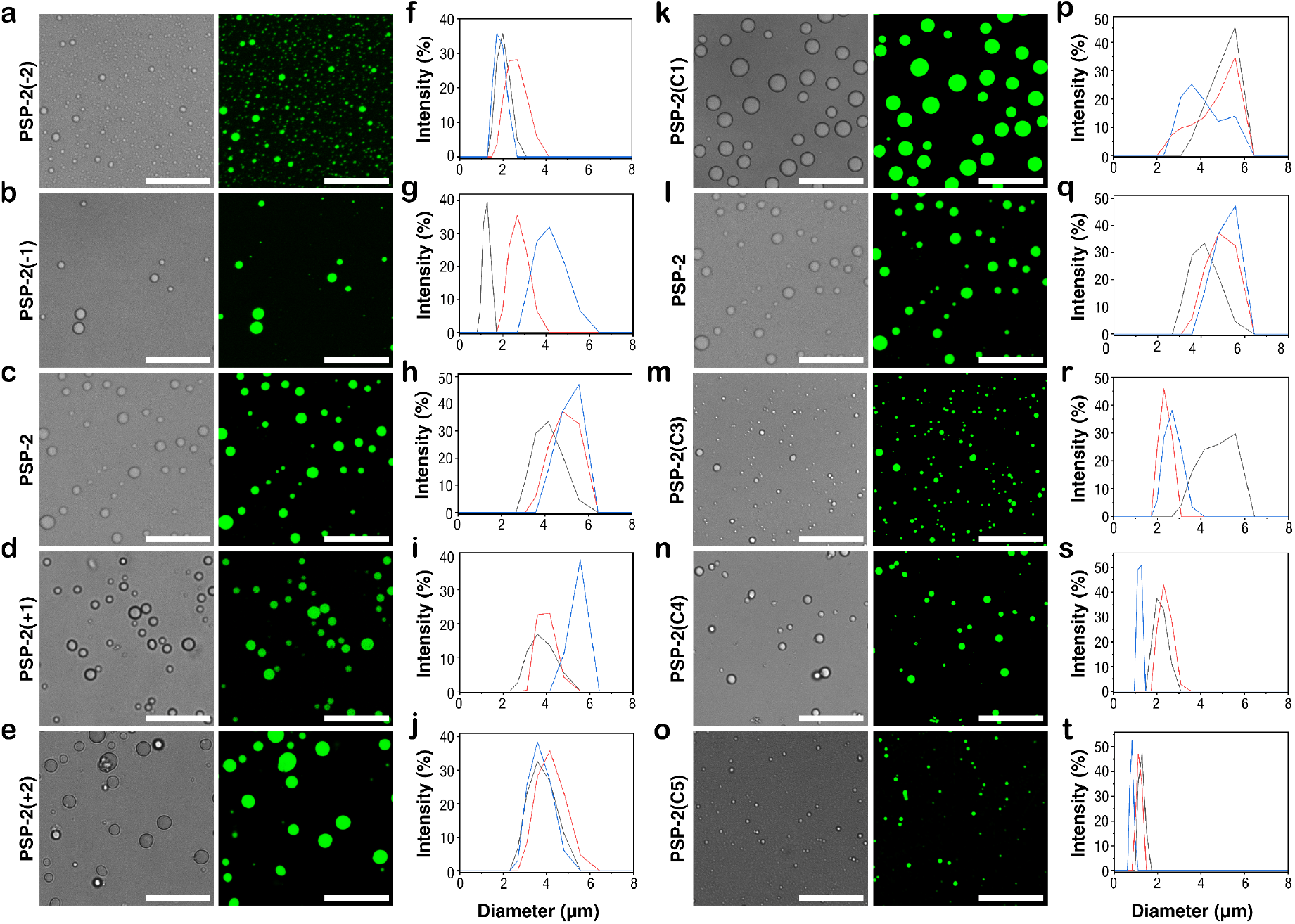
Morphology and size distribution of PSP condensates. (a-e and k-o): Representative bright-field and confocal images of condensates formed by spacer-length and cysteine density variants, respectively. Condensates were prepared in 20 mM Tris, 2.5 M NaCl at pH 8.0, and at approximately twice their experimentally determined *Csat* values (Figure 2). For confocal fluorescence imaging, samples contained 1% FITC-labelled peptide. (f-j and p-t): Condensate size distributions determined by dynamic light scattering (DLS) for the corresponding spacer-length and cysteine density variants, respectively. Scale bars represent 10 µm.

### Cysteine Density Alters Condensate Fluid Dynamics More Significantly Than Inter-Cysteine Lengths

To determine how peptide sequence modifications influence condensate fluid properties, especially viscosity and diffusion, we performed fluorescence recovery after photobleaching (FRAP) experiments on both spacer and cysteine variants of PSP-2 (Figure 4). While spacer length changes produced relatively modest shifts in condensate dynamics, cysteine density changes produced substantial effects on molecular mobility within the condensed phase. All condensates formed by the spacer variants displayed rapid FRAP recovery (Figure 4a) but showed a linear increase in the recovery rate constant (*k*) (reflecting the apparent diffusion coefficient, *D*) as a function of increasing spacer lengths from approximately 0.06 s⁻¹ for PSP-2(−2) to 0.14 s⁻¹ for PSP-2(+2) (Figure 4b). Similarly, the extent of FRAP recovery (i.e. percentage of fluorescence fraction recovered) increased linearly from approximately 77% to 92% with increasing spacer length (Figure 4c). These observations suggest that extending the spacer region reduces intermolecular constraints and facilitates molecular diffusion within the condensates.

**Figure 4.**
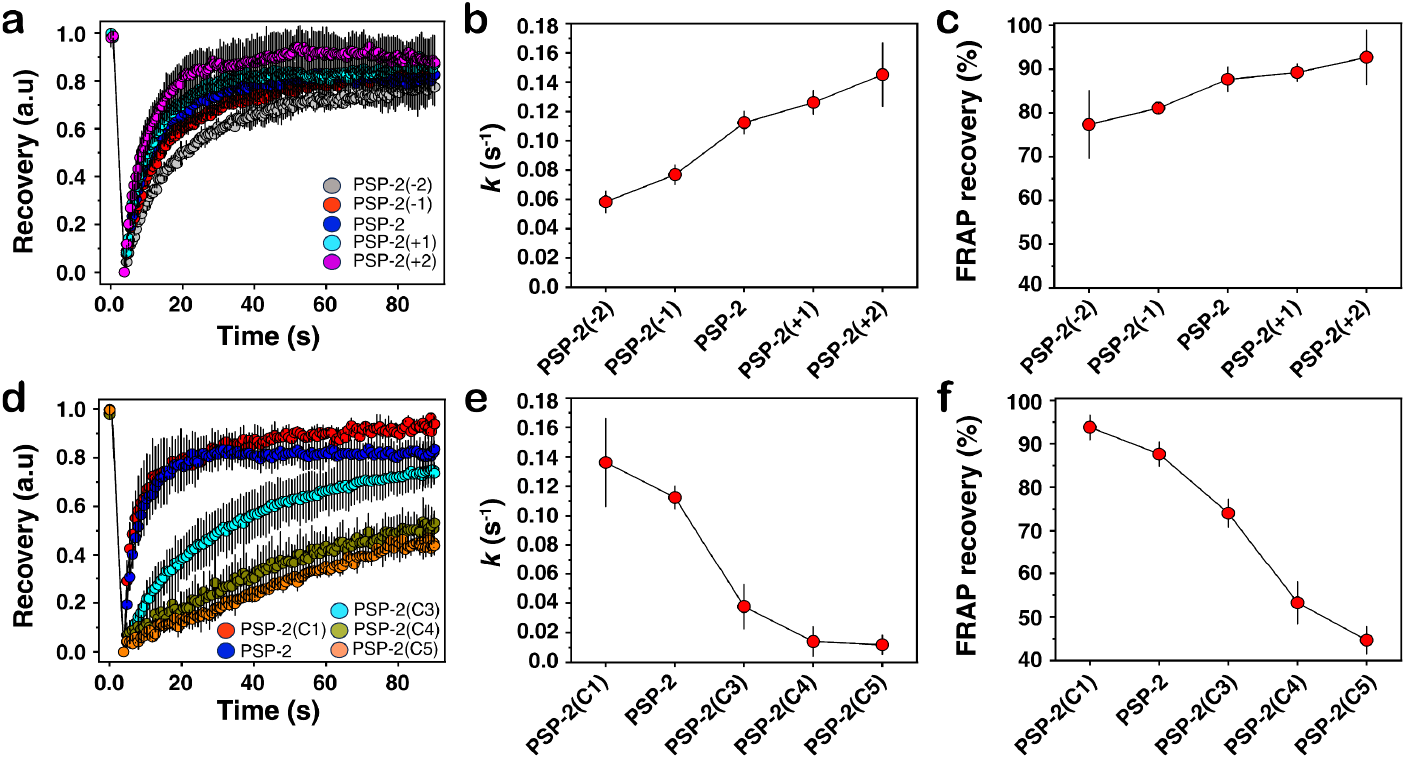
FRAP analyses of peptide condensates show distinctive differences between the two variant groups. (a) Temporal normalized FRAP kinetics for PSP spacer variants. (b) Corresponding recovery rate constants, *k,* derived from fitting the FRAP data with first-order exponential growth. (c) Percent fluorescence recovery of spacer variants. (d) Temporal normalized FRAP kinetics for PSP-2 cysteine variants. (e) Corresponding recovery rate constants, *k,* calculated from FRAP fitting of cysteine variants. (f) Percent fluorescence recovery of cysteine variants. (n = 3).

In contrast, varying cysteine density produced a much stronger effect on condensate dynamics (Figure 4d). The PSP-2(C1) and PSP-2(C2) variants exhibited rapid fluorescence recovery and high mobile fractions, with recovery values exceeding 85–90%, consistent with highly dynamic liquid-like condensates (red and blue; Figure 4d). However, increasing the number of cysteine residues progressively mitigated fluorescence recovery and the extent of FRAP recovery (Figure 4d). The recovery rate constant decreased by an order of magnitude, from approximately 0.14 s⁻¹ for PSP-2(C1) to 0.01 s⁻¹ for PSP-2(C5) (Figure 4e). Similarly, fluorescence recovery reduced from roughly 94% for PSP-2(C1) to only ∼45% for the PSP-2(C5) variant (Figure 4f), further indicating dramatically reduced diffusion within these condensates. These changes were much larger than those observed for the spacer variants, highlighting cysteine density as a dominant regulator of condensate viscosity.

The progressive loss of molecular mobility with increasing cysteine density is consistent with enhanced intermolecular disulfide-mediated interactions that generate a more interconnected network within the condensates. As a result, the condensates transition from highly dynamic, liquid-like assemblies to increasingly viscous, gel-like states. Collectively, these findings show that although spacer length can fine-tune condensate dynamics, cysteine density is a much stronger determinant of molecular mobility and internal condensate organization.

### Droplet Coalescence Reveals Condensate Fluidity

To better understand how spacer length and cysteine content affect condensate fluidity, we monitored changes in droplet coalescence over time. Representative time-lapse fluorescence images show that the different PSP variants differ clearly in their ability to fuse with droplets and relax back to a spherical shape (Figure 5a–j). We used changes in aspect ratio over time to compare shape-relaxation rates between variants.^24–27^ Among the spacer library, PSP-2 and PSP-2(−1) showed the fastest rates of relaxation (Figure 5a–e,k). In comparison, PSP-2(−2), PSP-2(+1), and PSP-2(+2) relaxed more slowly and maintained higher aspect ratios throughout the observation period, suggesting less fluidity. In particular, PSP-2(+2) showed the slowest change in shape, suggesting that increasing spacer length affects the condensates’ ability to fuse and reorganize. The cysteine variants showed a more pronounced difference (Figure 5f–j,l). PSP-2(C1) and PSP-2(C2) showed a rapid decrease in aspect ratio following droplet fusion. In contrast, PSP-2(C3), PSP-2(C4), and PSP-2(C5) showed only small changes and remained considerably elongated after contact or fusion on these time scales. This slower relaxation was more noticeable in variants with higher cysteine content. Together, these results show that both spacer length and cysteine density influence condensate coalescence. The faster relaxation observed for PSP-2, PSP-2(−1), and the lower-cysteine variants is consistent with more dynamic, liquid-like condensates, whereas the slower relaxation of PSP-2(+1), PSP-2(+2), and especially the higher-cysteine variants suggests more restricted molecular rearrangement within the condensates.

**Figure 5.**
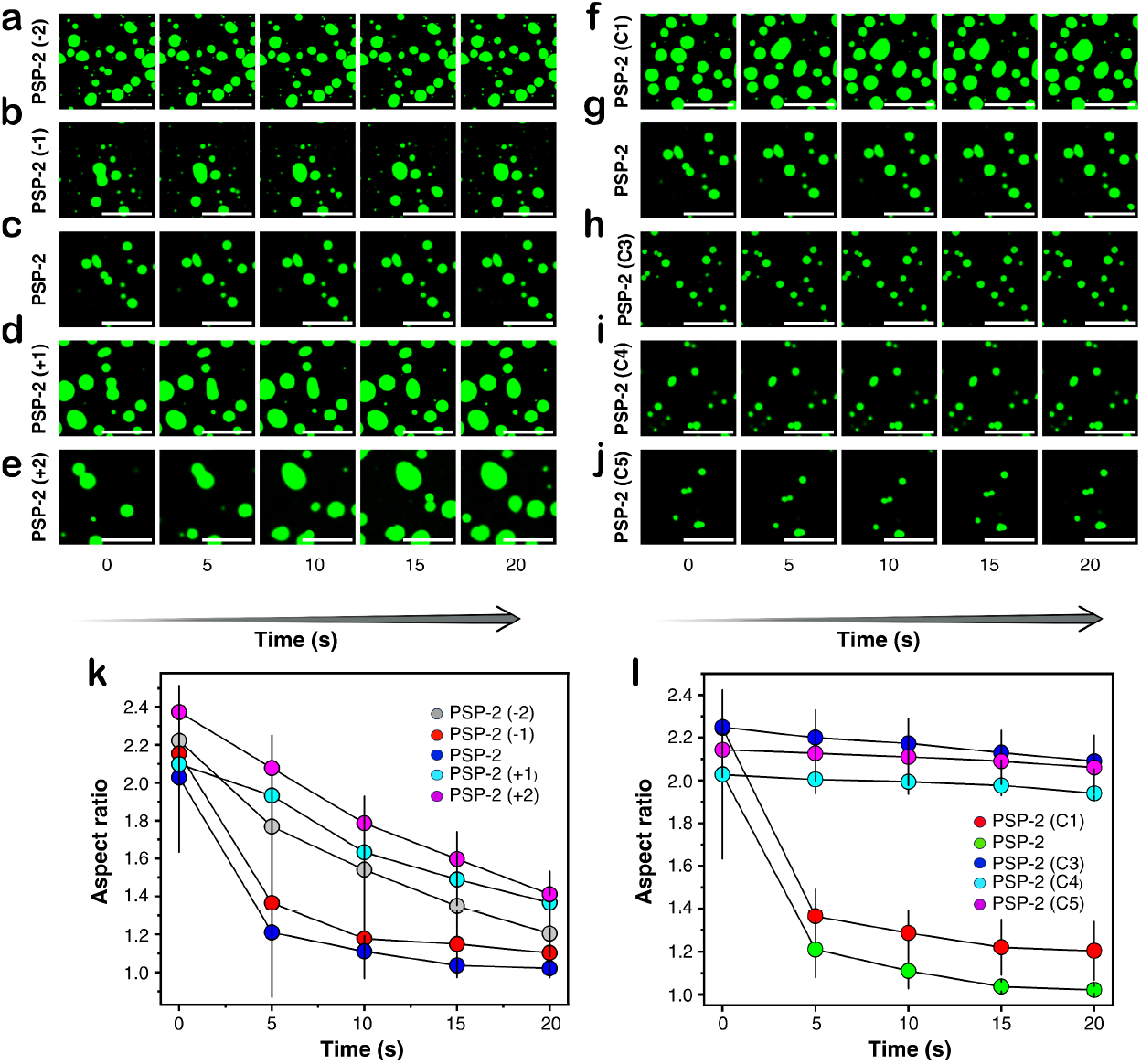
Droplet coalescence of PSP spacer and cysteine variants. Representative time-lapse images showing condensate fusion and shape relaxation for the PSP-2 spacer variants (a-e): PSP-2(−2), PSP-2(−1), PSP-2, PSP-2(+1), and PSP-2(+2), and the cysteine variants (f-j): PSP-2(C1), PSP-2(C2), PSP-2(C3), PSP-2(C4), and PSP-2(C5). For each variant, condensates were prepared at twice its respective saturation concentration (*Csat*), in 50 mM tris buffer containing 2.5 M NaCl at pH 8.0. One percent FITC-labeled peptide was included for fluorescence imaging. Scale bars represent 20 µm. (k, l) Changes in the aspect ratio of the spacer variants and cysteine variants, respectively, during the coalescence process. An aspect ratio approaching 1 indicates that the fused condensates became more spherical over time. Scale bars represent 20 µm.

### Condensate Maturation Suggests Greater Dependency on Cysteine Density Than Inter-Cysteine Lengths

Because biomolecular condensates transition into non-Newtonian, viscoelastic materials by percolation,^28^ we next examined how the condensates matured over longer incubation times. We observed the samples for up to 10 days and measured their dynamics by FRAP. PSP-2(−2), the variant with the shortest spacer, largely formed numerous small condensate droplets, with some large droplets appearing by day 10 (Figure 6a). PSP-2(−1), the second spacer variant with a slightly longer spacer, showed larger droplets at day 1 that became fewer by day 10, suggesting coalescence during incubation (Figure 6b). PSP-2(C2), PSP-2(+1), and PSP-2(+2), which have increasing spacer lengths, formed much larger condensates, but the droplet size and number differed considerably among the variants (Figure 6c–e).

**Figure 6.**
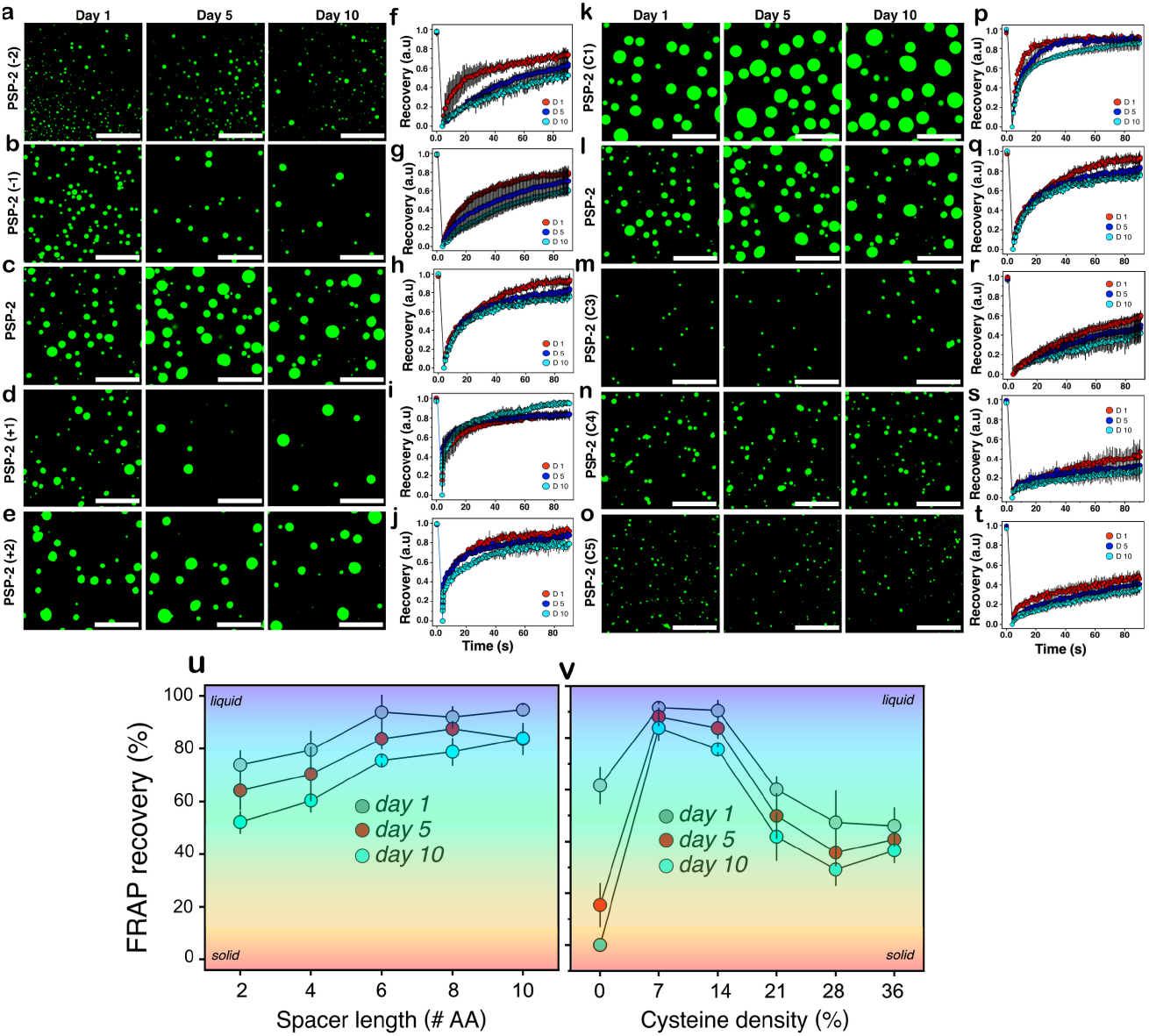
Temporal condensate maturation and dynamics. Confocal fluorescence microscopy images and FRAP analyses illustrating the aging-dependent evolution of condensates over 10 days of incubation. (a–e): Condensates imaged after days 1, 5, and 10 by the spacer variants, PSP-2(−2), PSP-2(−1), PSP-2(C2), PSP-2(+1), and PSP-2(+2). (f-j): Corresponding FRAP recovery curves (n >3). (k–o): Condensates imaged after days 1, 5, and 10 by the cysteine variants, PSP-2(C1), PSP-2(C2), PSP-2(C3), PSP-2(C4), and PSP-2(C5). (p-t): Corresponding FRAP recovery curves (n >3). (u-v): Compiled FRAP recovery plots as a function of spacer length and cysteine densities, respectively, overlaid with the approximate ranges of fluidity. Scale bars represent 20 µm.

The FRAP recovery curves for the spacer variants are shown in Figure 6f–j. The recovery rates and extent for peptides with short inter-cysteine lengths, PSP-2(−2) and PSP-2(−1), noticeably decreased from day 1 to day 10, suggesting possible percolation behavior that led to the formation of a gelated material (Figure 6f,g). Variants with longer inter-cysteine lengths, including PSP-2(C2), PSP-2(+1), and PSP-2(+2), showed higher recovery values than the shorter-spacer variants (Figure 6h–j and 5u). Variants with larger spacer lengths of 6–10 amino acids had a greater extent of FRAP recovery than those with shorter inter-cysteine lengths (Figure 6u). These data suggest that short inter-cysteine lengths may improve percolation by increasing sticker interactions through higher effective concentrations and promoting greater viscoelasticity compared with longer-spacer variants.

The cysteine density variants showed dramatically different behavior (Figures 6k–o). The control peptide without cysteines (PSP-2 (C0) has been established to form highly gelated material within three days of incubation.^21^ The one-cysteine-containing PSP-2(C1) formed large droplets that remained consistent over the 10-day incubation (Figure 6k). All cysteine variants showed an inverse correlation between droplet size and cysteine density, and for each peptide, the droplet size remained unchanged for most of the incubation period (Figure 6m–o). The FRAP recoveries of PSP-2(C1) and PSP-2(C2) were relatively high, suggesting dynamic condensates (Figure 6p,q). In contrast, PSP-2(C3), PSP-2(C4), and PSP-2(C5) recoveries were significantly blunted, suggesting low dynamics and diffusion (Figure 6r,s,t). Interestingly, FRAP recovery for cysteine-density variants showed non-monotonic behavior, unlike the spacer variants, with PSP-2(C1) and PSP-2(C2) showing maximum recovery, followed by decreasing recovery for PSP-2(C3), PSP-2(C4), and PSP-2(C5) variants (Figure 6v). This may be because higher cysteine content increases disulfide-crosslink density, making the condensates less fluid. Together, these results show that cysteine density affects condensate maturation more than the spacer lengths. Longer inter-cysteine lengths and intermediate cysteine densities promoted more fluid-like behavior, while short inter-cysteine lengths or high cysteine densities resulted in gel-like condensate behavior. If one can approximate material properties from FRAP recovery, the data suggest that varying cysteine density can control the material properties of the condensates formed (Figure 6v).

### Cysteine Density Affects Redox Reversibility of Condensate Formation and Dissolution

We previously showed that PSP-2 condensates reversibly dissolve under reducing conditions and reform under oxidizing conditions.^21^ To test whether the peptide condensates could dissolve and reform across redox cycles, we exposed the spacer and cysteine variants to three cycles of reduction and oxidation (Figure 7). We prepared the peptide samples at twice their respective saturation concentration (*C_sat_*) in 20 mM Tris buffer containing 2.5 M NaCl at pH 8.0. The turbidity plots for the spacer variants show that turbidity decreased immediately upon DTT addition, indicating condensate dissolution (first blue data point; Figure 7a, c, e, g, and i). Corresponding fluorescence microscopy images confirm the dissolution, as no droplets were visible under reducing conditions (images in blue squares). Adding H₂O₂ caused a sharp increase in turbidity and the corresponding reappearance of droplets (second red data point; Figures 7a, c, e, g, and i). This dissolution-and-reformation behavior was observed unabated for two more cycles. The FRAP recovery data before and after reduction and oxidation confirm the reversibility with nearly identical recovery curves (Figure 7(b, d, f, h, and j). However, the cysteine variants showed variability; reversibility depended on cysteine density (Figure 7k–r). PSP-2(C1), PSP-2(C2), and PSP-2(C3) showed reversibility, with a sharp decrease in turbidity upon DTT treatment, followed by an increase upon H₂O₂ oxidation (Figure 7k, m, and e). Although C1 showed a steady decrease in turbidity values after each cycle (Figure 7k), both confocal images and FRAP recovery curves confirm reversibility (Figure 7k and l). In contrast, PSP-2(C4) and PSP-2(C5) did not fully dissolve under reducing conditions (Figures 7o and q). Although turbidity decreased slightly after DTT treatment, the values remained well above baseline, and droplets were still visible in the fluorescence images (Figures 7o and q). The corresponding FRAP recovery curves for all the cysteine variants also correlated with the turbidity trends (Figures 7l, n, p, and r). The data show that condensates formed by peptides with high cysteine density (PSP-2(C4) and PSP-2(C5)) resist dissolution in a reducing environment and contrast sharply with other variants, which show redox reversibility over three cycles.

**Figure 7.**
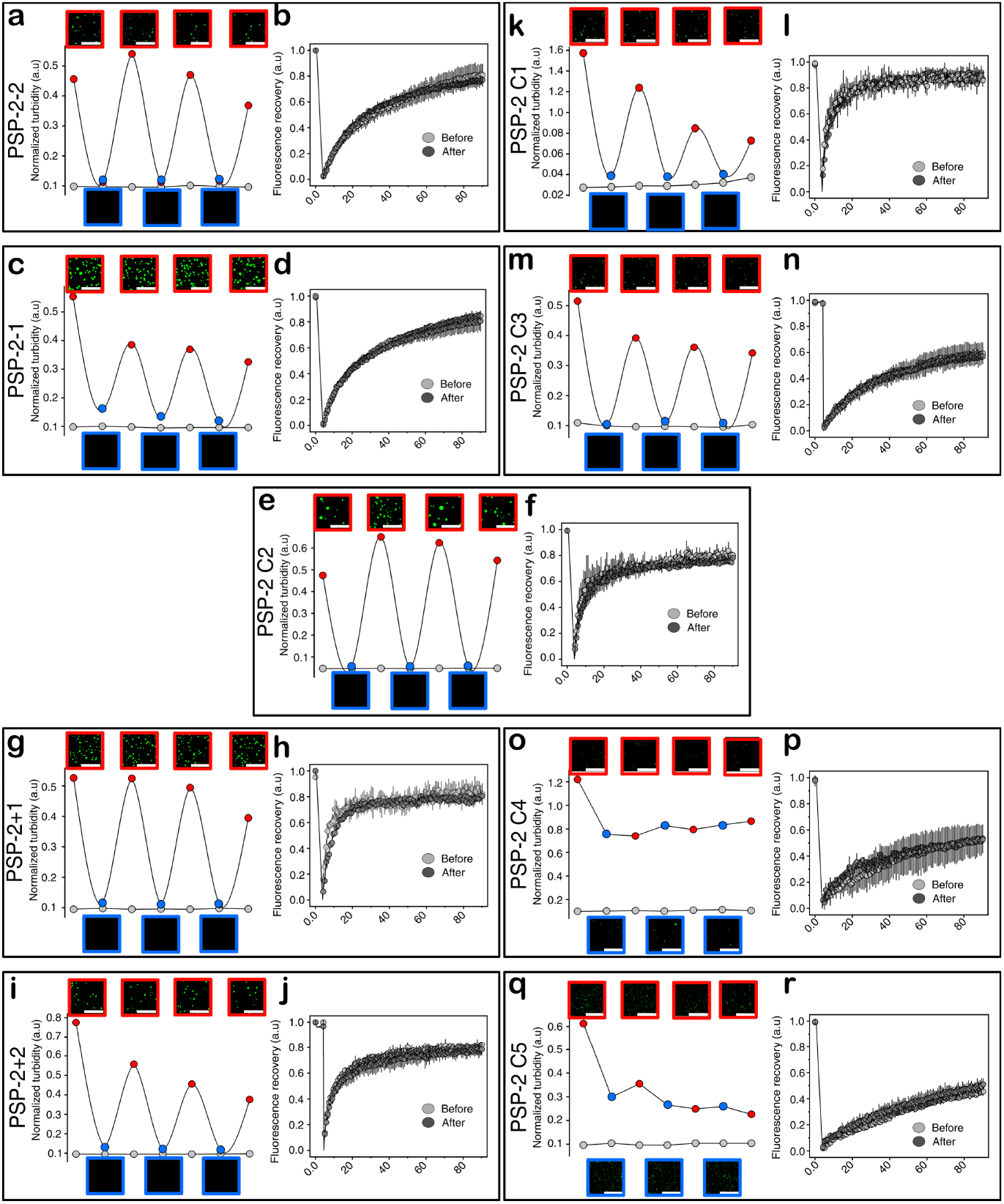
Multi-cycle reversibility of condensate formation and dissolution under redox flux. Normalized turbidity measurements of PSP spacer and cysteine variants under reducing (blue circles) and oxidizing (red circles) conditions. Peptide concentrations were kept at twice their respective saturation concentrations (*Csat*) in 20 mM Tris, 2.5 M NaCl, and pH 8.0. Turbidity was monitored at 600 nm during sequential reduction (DTT) and reoxidation (H₂O₂) cycles. In the first cycle, a five-fold molar excess of DTT relative to the number of cysteines was added to ensure complete reduction, followed by a sequential increase in subsequent cycles. Color-coded fluorescence microscopy images for each redox cycle are shown above and below the turbidity traces. FRAP recovery profiles before the first reduction (grey circle) and after the last reduction (black circle). Panels a-j correspond to spacer variants, and k-r correspond to the cysteine variants. (n = 3). Scale bars represent 20 µm.

Kinetics of condensate dissolution in the presence of a reducing agent also reveal how spacer and cysteine density influence disulfide-bond resistance. In parallel, we incubated the samples in Figure 7 with a five-fold molar excess of DTT relative to cysteines and monitored dissolution kinetics by turbidity (Figure 8). The data showed that the dissolution rate increased linearly with spacer length (Figure 8a) and that, for the cysteine variants, dissolution rate decreased with cysteine density (Figure 8b). Among the cysteine variants, PSP-2(C5) showed the slowest rate, further confirming that disulfide bonds resist reduction and hence PSP dissolution in high-cysteine-density condensates.

**Figure 8.**
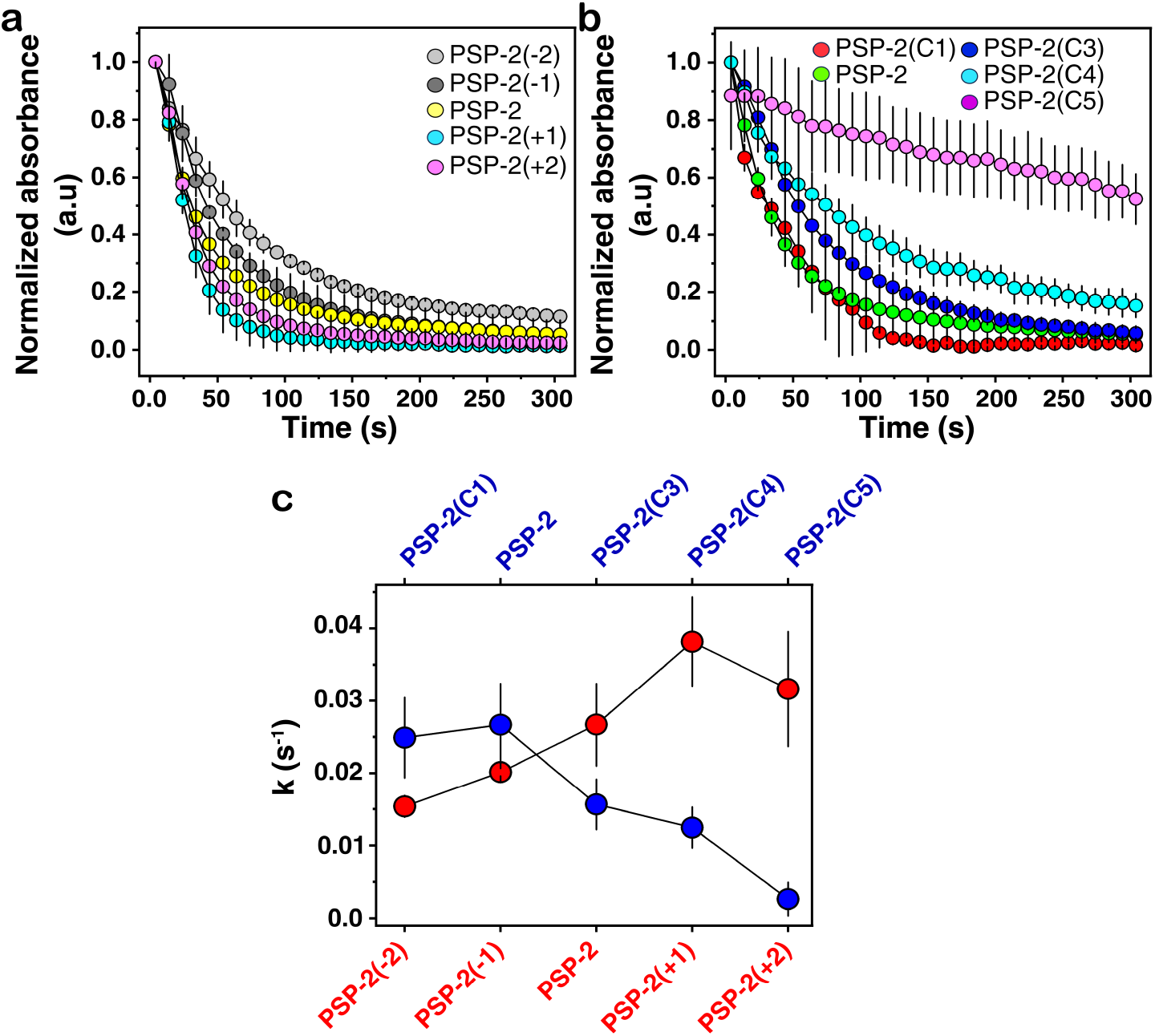
Dissolution kinetics of spacer and cysteine variants. a and b: Baseline-corrected and normalized temporal turbidity measured as absorbance at 600 nm for PSP spacer and cysteine variants, respectively, in the presence of the reducing agent DTT. Peptides were incubated for 30 minutes at twice their respective *Csat* in 20mM Tris, 2.5 M NaCl (pH 8.0) to allow droplet formation. DTT was then added at a 5-fold molar excess relative to the total cysteine content, and then turbidity measurements at 600nm were initiated immediately to monitor droplet dissolution. C: Rate constants (*k*) obtained by fitting the kinetic data with a first-order exponential decay model.

### Encapsulations of a Protein and Small Molecule Provide Clues on Condensate Mesh Size

Because disulfide crosslink density directly affects condensate pore size, we assessed pore size in the cysteine variants by estimating encapsulation efficiency (EE) and diffusion of a large cargo (eGFP) and a small molecule (tetramethylrhodamine, TMR). eGFP was clearly enriched inside the condensates formed by PSP-2(C2). However, the fluorescence intensity inside the condensates gradually decreased for PSP-2(C3), PSP-2(C4), and PSP-2(C5). Interestingly, PSP-2(C4) and PSP-2(C5) variants localized on the periphery of the condensates (Figure 9a). EE decreased with increasing cysteine density, with PSP-2(C2) highest (∼30%) and PSP-2(C5) lowest (∼5%) (Figure 9c). A similar trend was observed for TMR, although the decrease in encapsulation was less pronounced than with eGFP; PSP-2(C2) showed the highest %EE (∼60%), and PSP-2(C3), PSP-2(C4), and PSP-2(C5) showed gradually decreasing EEs. (Figure 9b and d). Unfortunately, %EE of TMR in PSP-2(C1) could not be determined because of TMR precipitation due to high concentrations needed to maintain the constant peptide-to-dye ratio. Nevertheless, both cargo showed similar decreasing trends with increasing cysteine density, with TMR showing higher EE than eGFP, as expected due to their sizes (Figure 9c and d). These decreasing trends reflect the effect of a smaller mesh size due to higher disulfide crosslink density.

**Figure 9.**
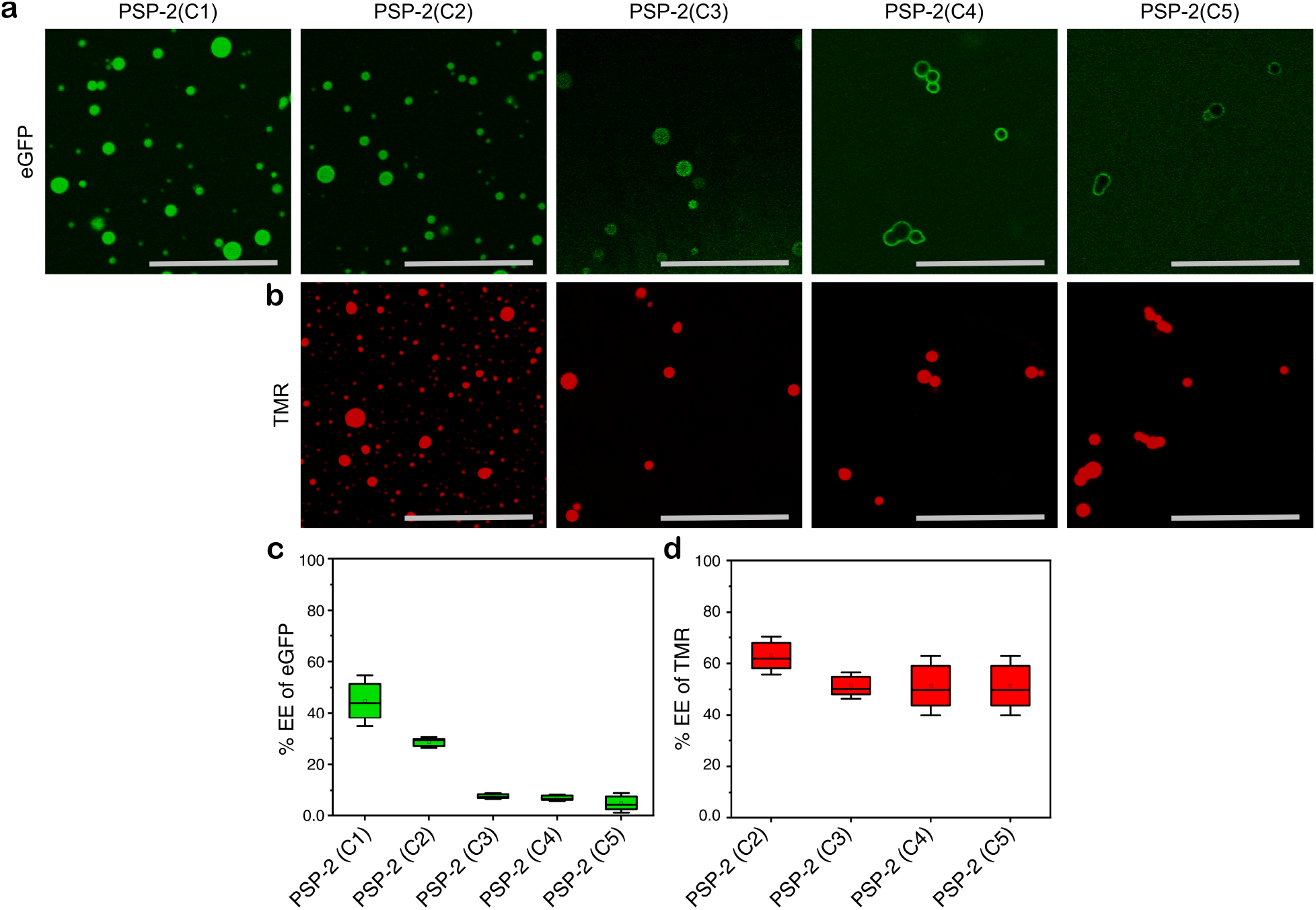
Encapsulation of eGFP and TMR in cysteine variants. Fluorescence microscopy images showing the encapsulation of eGFP (a) and TMR (b) within PSP-2 cysteine variants, PSP-2(C1) to PSP-2(C5). The images show that the amount of encapsulated cargo changes across the different cysteine variants. (c) Quantification of the encapsulation efficiency (% EE) of eGFP for each variant. (d) Quantification of the encapsulation efficiency (% EE) of TMR. Overall, encapsulation efficiency gradually decreases from PSP-2(C1) to PSP-2(C5), with a stronger decrease for eGFP than for TMR.

## DISCUSSION

Biomolecular condensates are now well recognized as important viscoelastic, membraneless hubs responsible for many cellular processes across all kingdoms of life. While condensate formation is driven by weak, multivalent non-covalent interactions (such as π–π or cation–π interactions),^16,20,23,30^ interspersing cysteines within disordered protein/peptide sequences introduces a covalent dimension in condensate design and biology. We explored this dimension and established key guiding principles.^21,22^ Under oxidizing conditions, disulfide bonds act as tunable crosslinks that lower the critical saturation concentration (*C_sat_*) for phase separation, facilitating stress-triggered condensate assembly.^21^ We also observed that 14% cysteine density (two cysteines) forms condensates that maintain their fluidity for prolonged periods, possibly inhibiting percolation toward viscoelastic behavior.^21^ The degree of disulfide crosslinking appears to directly control condensate viscoelasticity, as we and others have shown.^21,31,32^ In this work, we focused on gaining detailed insights into how cysteine density and inter-cysteine distance (spacer length) affect condensate properties by designing peptides that systematically vary both parameters (Figure 1).

The data presented here show that spacer length and cysteine density affect condensate viscosity and maturation. Collectively, our results show that cysteine density imparts a greater influence on condensate properties than the spacer length. Spacer lengths affect cation–π interactions between R and Y through entropic penalties, thereby changing the effective concentrations of the interacting nodes (R and Y), which inversely correlate with length, as shown in the *C_sat_* plots (Figure 2). Subtle yet tangible effects of inter-cysteine length are also evident in coalescence behavior and dynamics (Figures 3 and 4), maturation (Figure 5), and dissolution kinetics (Figure 6). As spacer length increases, cysteine density also changes in these variants, ranging from 10% (PSP-2(−2)) to 18% (PSP-2(+2)). However, only amino acids S and G vary; they are expected to be benign because they do not interact with R’s, Y’s, or cysteines and are therefore assumed not to contribute to sticker interactions. In sharp contrast, cysteine density strongly affects condensate properties. First, we vary cysteines from 0 to 35% while keeping the spacer length (S and G) constant (two residues). Although the inter-cysteine distance is more than two spacers (S and G), the other residues are arginine, which are expected to be involved in multivalent interactions. Increasing cysteine density sharply decreased condensate *C_sat_* (Figure 2) and dramatically reduced fluidity and coalescence, especially above 21% (C3-C5) (Figures 3-6). Interestingly, cysteine density increases non-monotonically: up to 14% density maintains fluidity, while 21-35% exhibit increased viscoelasticity (Figure 6). Importantly, condensates with high cysteine densities showed no redox reversibility and decreased cargo encapsulation efficiency, suggesting that high disulfide-bond crosslinking can serve as a tunable lever to control material state, redox sensitivity, and condensate porosity. This likely relates to cysteines forming disulfide bonds, which provide additional connections between peptide molecules. As more cysteines are introduced, the number of possible intermolecular crosslinks also increases, making it easier to form and stabilize a peptide-rich network. The much larger change in *C_sat_* across the cysteine series than across the spacer series suggests that cysteine content more strongly regulates phase separation in this system.

This work also raises several important questions. One is whether cysteines, through disulfide crosslinks, can promote condensate formation on their own, without weak non-covalent interactions between R and Y. Our investigations used a control peptide lacking canonical interacting nodes, R’s and Y’s (C-S-G-S-G-G-G-S-G-S-C), a variant of PSP-2(−2) that failed to show phase separation even up to 10 mM concentrations under the same buffer conditions as the other peptides (Figure S3). This suggests that π–π or cation–π interactions are required for phase separation and that cysteines could act as auxiliary interaction nodes, as we previously concluded.^21^ The second question concerns the observation that condensates with high cysteine density remain perpetually “arrested” in contact and do not coalesce, unlike other droplets (Figure 4i and j; Figure S4a). We speculate that dense disulfide crosslinks within the condensates increase interfacial tension and drive this phenomenon. In addition, as mass spectrometry shows (Figure S5), some free thiols on the droplet surface may form disulfide bonds with cysteines from neighboring droplets, crosslinking the already strained droplets (due to interfacial tension) and arresting coalescence (Figure S4a and b). Lastly, it is unclear why eGFP fails to partition into condensates with high cysteine density (C4 and C5) and instead remains on their surface (Figure 9). Although pore size is an important factor, we speculate that, in conjunction with the interfacial tension imparted by the high density of cysteine disulfide crosslinks mentioned above, the peptides are highly positively charged due to arginine inclusion. This could also localize eGFP on the surface, which has a net negative charge at pH 8.0, accentuated by the pore-size limits (Figure S5c and d).

## CONCLUSION

In recent years, biomolecular condensates have emerged as attractive scaffolds for soft-material design in various biotechnological applications, as their viscoelasticity can be modulated to tailor tissue-specific cell proliferation and clearance.^33–36^. Systematically modulating condensate fluidity through cysteine incorporation provides an effective strategy for controlling the material properties of peptide-based biomolecular condensates. This moves beyond the traditional “stickers and spacers” model^16,23^ by providing a chemically specific, redox-responsive mechanism to tune condensate stiffness, dynamics, and stability—making cysteine-rich disordered proteins/peptides a biologically important class of molecules. Beyond fundamental biology, rationally engineering condensates whose assembly, viscoelasticity, and porosity can be modulated by redox signals has transformative implications for biotechnology and biomaterials design. Stimuli-responsive materials that sense and respond to chemical environments are among the most sought-after targets in materials science, yet achieving this goal remains elusive, especially at the nanoscale with molecular precision. More broadly, the established design principles will provide the field with a generalizable engineering toolkit for constructing synthetic condensates, responsive hydrogels, and tunable bioreactors for applications in drug delivery, synthetic biology, and regenerative medicine.

## AUTHOR INFORMATION

Conceptualization – VR; intellectual inputs – VR, MM, TDC, CTA, WSS; biophysical experiments, data interpretation – MM, VR, CTA, TDC; peptide synthesis – WSS, HL, TDC; data processing – MM, HL, CTA; manuscript writing and editing – VR, MM, TDC.

## ACKNOWLEDGEMENTS

The authors thank the National Science Foundation (BMAT 2208349) and (ART 2331378) for each providing 50% financial support for this project. The authors also acknowledge support from the National Science Foundation (Award number: 2319932) for mass spectrometry instrumentation used in this study. TDC acknowledges funding support from the NIH and the National Institute of Biomedical Imaging and Bioengineering (NIBIB R21EB033533). WSS gratefully acknowledges fellowship support from the Mississippi Space Grant Consortium (MSSGC) funded by the National Aeronautics and Space Administration (NASA) Office of STEM Engagement.

## SUPPLEMENTARY INFORMATION

### MATERIALS AND METHODS

#### Materials

Acetonitrile, *N*,*N* -dimethylformamide (DMF), diethyl ether, 2,2’-(ethylenedioxy)diethanethiol (EDT), and 4-methylpiperidine were all purchased from Sigma-Aldrich and used as received. All Fmoc-protected amino acids, Oxyma, and rink amide MBHA resin were purchased from CEM. *N*,*N* -diisopropylcarbodiimide (DIC) was purchased from AA Blocks and used as received. Trifluoroacetic acid (TFA) was purchased from Toyko Chemical Industry and used as received. Triisopropylsilane (TIPS) was purchased from Chem Impex International and used as received.

#### Peptide Synthesis

Peptides were synthesized using Fmoc-solid-phase peptide chemistry on a CEM Liberty Blue microwave-assisted peptide synthesizer (CEM, Matthews, NC, USA). Fmoc groups were cleaved using 20% 4-methylpiperidine in *N*,*N*-dimethylformamide (DMF) at 90 °C for 30 s. Amino acids were coupled using 4 mol equivalents of each Fmoc-protected amino acid, 8 equivalents ethyl cyanohydroxyiminoacetate (Oxyma) and 8 equiv of *N*,*N*′-diisopropylcarbodiimide (DIC) for 2–4 min at 90 °C in DMF as solvent. Completed peptides were then cleaved from the resin using a solution of 92.5:2.5:2.5:2.5 trifluoroacetic acid (TFA)/triisopropylsilane (TIPS)/ 2,2’-(ethylenedioxy)diethanethiol (EDT) /DI water for 4-5 h at ambient temperature on a wrist shaker. Following cleavage, peptides were precipitated with cold diethyl ether, collected via centrifugation, and dried overnight. The peptides were then purified by preparative-scale reverse-phase high-performance liquid chromatography (CEM Prodigy HPLC) using a Phenomenex Gemini column (C-18 stationary phase, 5 μm, 100 Å pore size, 50 × 250 mm). The mobile phase was acetonitrile and water, both containing 0.1% TFA. Pure fractions were identified using electrospray ionization mass spectrometry (ESI-MS) in positive ion mode on a Thermo Scientific Orbitrap Exploris™ 240 mass spectrometer using direct injection. Excess acetonitrile was removed by rotary evaporation, the samples were freeze-dried, and the powders were stored at −20 °C until use.

#### Preparation of Condensates

Based on the *C_sat_* estimated in turbidity assay and sedimentation assay for each peptide self-coacervation, the lyophilized powder of the HPLC-purified peptide derivatives was dissolved in autoclaved, nanopure water, and the stock solution of ∼ 20-30 mM was stored at −20 °C until further use. Prior to the experiments, an aliquot of the peptide stock was taken to make a self-coacervate sample of concentrations twice their respective *C_sat_*; i.e., 600 µM of PSP-2(−2), 1mM of PSP-2(−1), 1.5 mM of PSP-2, 1 mM of PSP-2(+1), 2 mM of PSP-2(+2), PSP-2(C3) and PSP-2(C4) in 50 mM Tris buffer at pH 8.0 with 2.5 molar NaCl and incubated for around 15 minutes at RT.

#### Phase Diagrams

Phase diagrams for peptide self-coacervation were constructed using a combination of turbidity measurements and centrifugation-based depletion assays. Peptide variants {PSP-2(−2), PSP-2(−1), PSP-2, PSP-2(+1), PSP-2(+2), PSP-2(C3), and PSP-2(C4)} were prepared across a range of concentrations with 2.5 M NaCl under buffered conditions (pH 8.0).

#### Turbidity assay

A BioTek Synergy H1 microplate reader was used to measure turbidity. Before each measurement, reactions were started and allowed about ten minutes to equilibrate at room temperature. Stocks were made at high quantities to avoid sample dilution for the peptide and NaCl tests. For samples where liquid droplets have been verified by optical microscopy, we employ turbidity as an indicator of LLPS. A boundary value of 1.30 A.U. optical density at 600 nm was used to create phase diagrams, and readings above this value were regarded as being in a phase transition. All measurements were made at room temperature. On Origin 8.5, data processing was carried out. We determined the mean from at least three datasets.

#### Centrifugation assay

To experimentally further confirm the saturation concentration for phase separation of each peptide, we performed a centrifugation-based depletion assay guided by turbidity measurements. A range of peptide concentrations was selected based on the phase boundaries identified from turbidity analysis. For each concentration, two parallel conditions were prepared: (i) a phase-separating condition and (ii) a non–phase-separating condition serving as a control. Samples were incubated at room temperature for 20 minutes to allow equilibration and potential condensate formation. Following incubation, samples were centrifuged at 12,000 rpm for 10 minutes to pellet any phase-separated material. The supernatant was collected carefully, and soluble peptide concentration was quantified by UV absorbance at 280 nm. The *C_sat_* was defined as the lowest total peptide concentration at which a measurable depletion of peptide in the supernatant was observed in the phase-separating conditions relative to the non–phase-separating control. This depletion indicates peptide partitioning into the condensed phase.

#### Dynamic Light Scattering (DLS)

The hydrodynamic size distributions of PSP condensates were characterized using dynamic light scattering (DLS). Condensates were prepared in 20 mM Tris buffer containing 2.5 M NaCl at pH 8.0, at approximately twice their experimentally determined saturation concentrations (Csat). Before measurements, the samples were diluted twofold using the same buffer to obtain suitable concentrations for DLS analysis. The diluted samples were then transferred to appropriate measurement cuvettes, and their hydrodynamic size distributions were determined using a dynamic light scattering instrument.

#### Reversible oxidation and reduction of the coacervates

Reversible peptide coacervation driven by redox modulation of disulfide bonds was investigated using combined turbidity measurements and confocal fluorescence microscopy. Peptide samples (700 µL total volume) were prepared at concentrations corresponding to twice their respective saturation concentration in 50 mM Tris buffer (pH 8.0) containing 2.5 M NaCl. For each cycle, a 100 µL aliquot was first withdrawn to measure turbidity at 600 nm using a BioTek plate reader. Subsequently, a 50 µL portion of the same sample was mixed with 0.5% FITC-labeled peptide and transferred onto a glass slide for imaging to capture initial coacervate formation using confocal fluorescence microscopy. To induce reduction, dithiothreitol (DTT) was added to the remaining reaction mixture at a 5-fold molar excess relative to cysteine residues. After 5 minutes of incubation, turbidity measurements and fluorescence imaging were repeated as described above. Reoxidation was then achieved by adding hydrogen peroxide (H₂O₂) at a 5-fold molar excess, followed by the same measurement and imaging procedures. This reduction–oxidation cycle was repeated four times to assess the reversibility and stability of peptide coacervation.

#### Coating glass slides and coverslips

Microscopic glass slides and coverslips were cleaned with 70% ethanol by sonicating for 15 minutes. Coverslips and glass slides were then allowed to air dry. 30 minutes were spent with the glass slides and coverslips were submerged in the coating solution (20% Tween20). To remove excess coating solution, the glass slides and coverslips were rinsed eight to ten times with Milli-Q water. Overnight, the glass slides and coverslips were dried at 37°C. Lens paper was used to wrap dry coated glass slides and coverslips, which were then kept at room temperature until needed.

#### Confocal microscopy and FRAP analysis

Confocal fluorescence imaging of peptide coacervates, including both self-coacervates and complex coacervates, was performed using a Leica STELLARIS DMi8 microscope equipped with a 63× objective. Samples were prepared on coated cavity glass slides, and droplets were allowed to settle for a few minutes prior to imaging to ensure stability. For fluorescence visualization, 0.5% FITC-labeled peptide was added to the reaction mixture after droplet formation. The internal dynamics of the condensates were characterized using fluorescence recovery after photobleaching (FRAP). Selected droplets were photobleached by applying a high-intensity laser (∼90%) for 5 seconds. Following photobleaching, fluorescence recovery within the bleached region was monitored over a 60-second period. Fluorescence recovery data were normalized and analyzed as a function of time using Origin 8.5 to evaluate the dynamic properties of the condensates.

#### Droplet Coalescence and Diffusion Analysis

Condensates of the PSP-2 spacer and cysteine variants were prepared using each peptide at twice its respective saturation concentration (*C_sat_*) in 50 mM Tris buffer containing 2.5 M NaCl at pH 8.0. For fluorescence imaging, 1% FITC-labeled peptide was mixed with the corresponding unlabeled peptide. A small volume of each condensate sample was placed on a clean, coated cavity glass slide, and droplet fusion events were recorded using a Leica STELLARIS-DMI8 microscope at 63× magnification. The recorded images were analyzed using Fiji (ImageJ). An ellipse was fitted around the boundary of each pair of fusing condensates, and the aspect ratio was calculated by dividing the major axis by the minor axis:

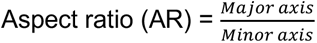

The decrease in aspect ratio over time was used to follow the relaxation of the fused condensates toward a spherical shape. An aspect ratio close to 1 indicated that the condensate had become nearly spherical. To maintain consistency among the different peptide variants, only fusion events involving condensates of approximately similar sizes were included in the analysis.

Where *r* is the radius of the bleached zone.

#### Dye Partitioning and Encapsulation

Peptide solutions were oxidized and diluted as described above. To each solution, 5 µM of dye was added and incubated for 10 minutes. Standard solutions for each equivalent dye and buffer concentration were prepared, and their UV-vis absorbances were recorded. The peptides were centrifuged at 15,000 × g for 10 minutes. Of the 100 µL volume, 40 µL was drawn as the supernatant, and the UV-vis absorbance was measured. The encapsulation efficiency of the dye was calculated using the following equation, where *A_sup_* is the absorbance of the supernatant, and *A_tot_* is the total absorbance at 490 and 550 nm for fluorescein and TMR, respectively.

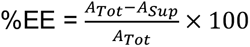

#### MALDI-ToF mass spectrometry

MALDI-TOF mass spectrometry was used to characterize IA-capped C5 peptide species under LLPS conditions. Condensates were formed in 50 mM Tris buffer, pH 8.0, containing 2.5 M NaCl. After condensate formation, the bulk sample was divided into two portions. One portion of the bulk condensate-containing solution was directly incubated with excess iodoacetamide (IA) to cap accessible cysteine residues and was analyzed as the total sample. A second portion of the same bulk sample was centrifuged to separate the condensate phase from the surrounding solution. The resulting supernatant was collected and incubated with excess IA for 1 h. Both the total sample and the supernatant fraction were analyzed by MALDI-TOF mass spectrometry using a Bruker Daltonics Microflex LT/SH TOF-MS system. Peaks corresponding to +3IA, +4IA, and +5IA were assigned to peptide species containing three, four, or five IA-capped cysteine residues, respectively.

### SUPPLEMENTARY FIGURES

**Figure S1.**
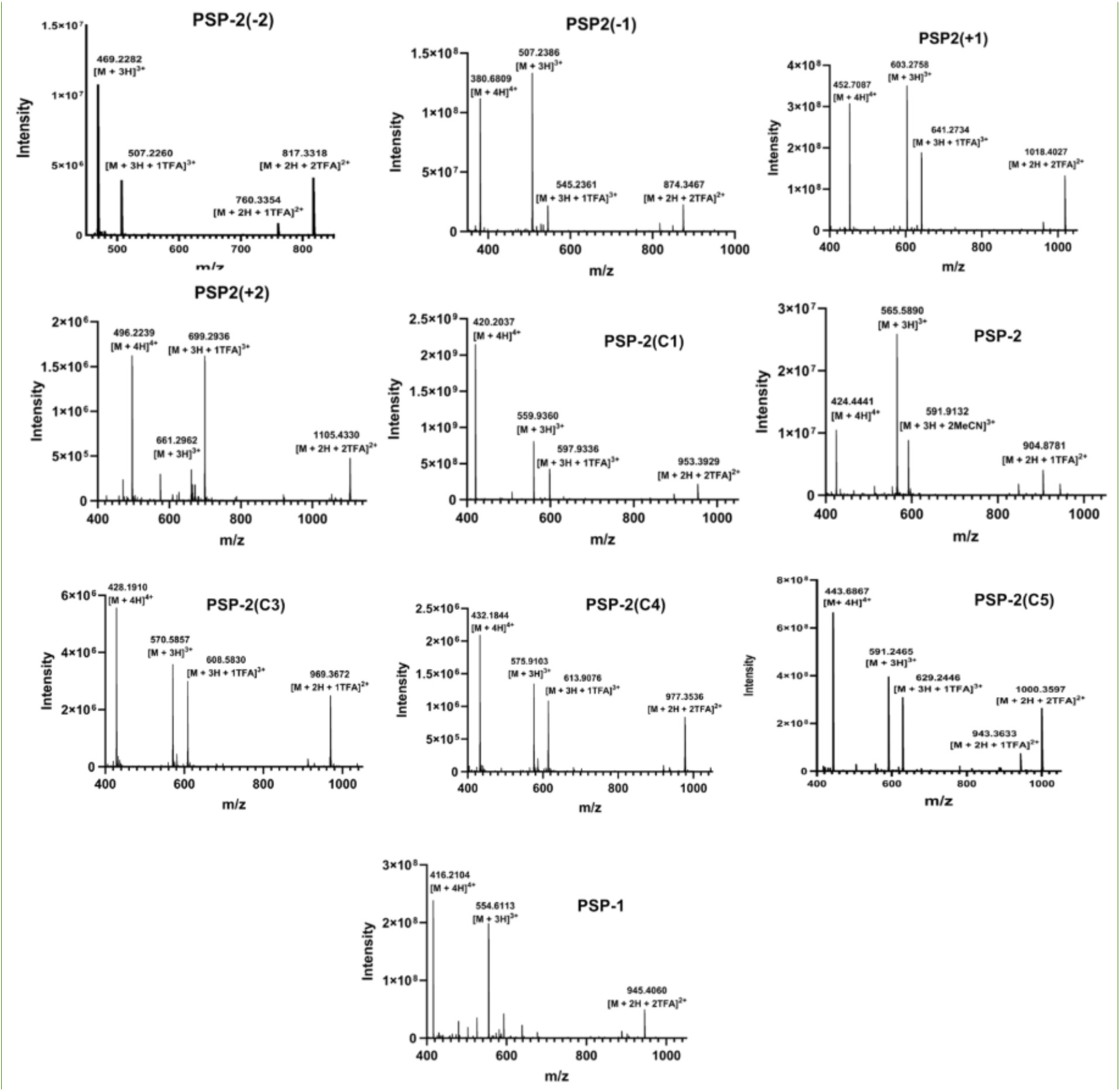
Purity of synthetic peptides confirmed by ESI mass spectrometry.

**Figure S2.**
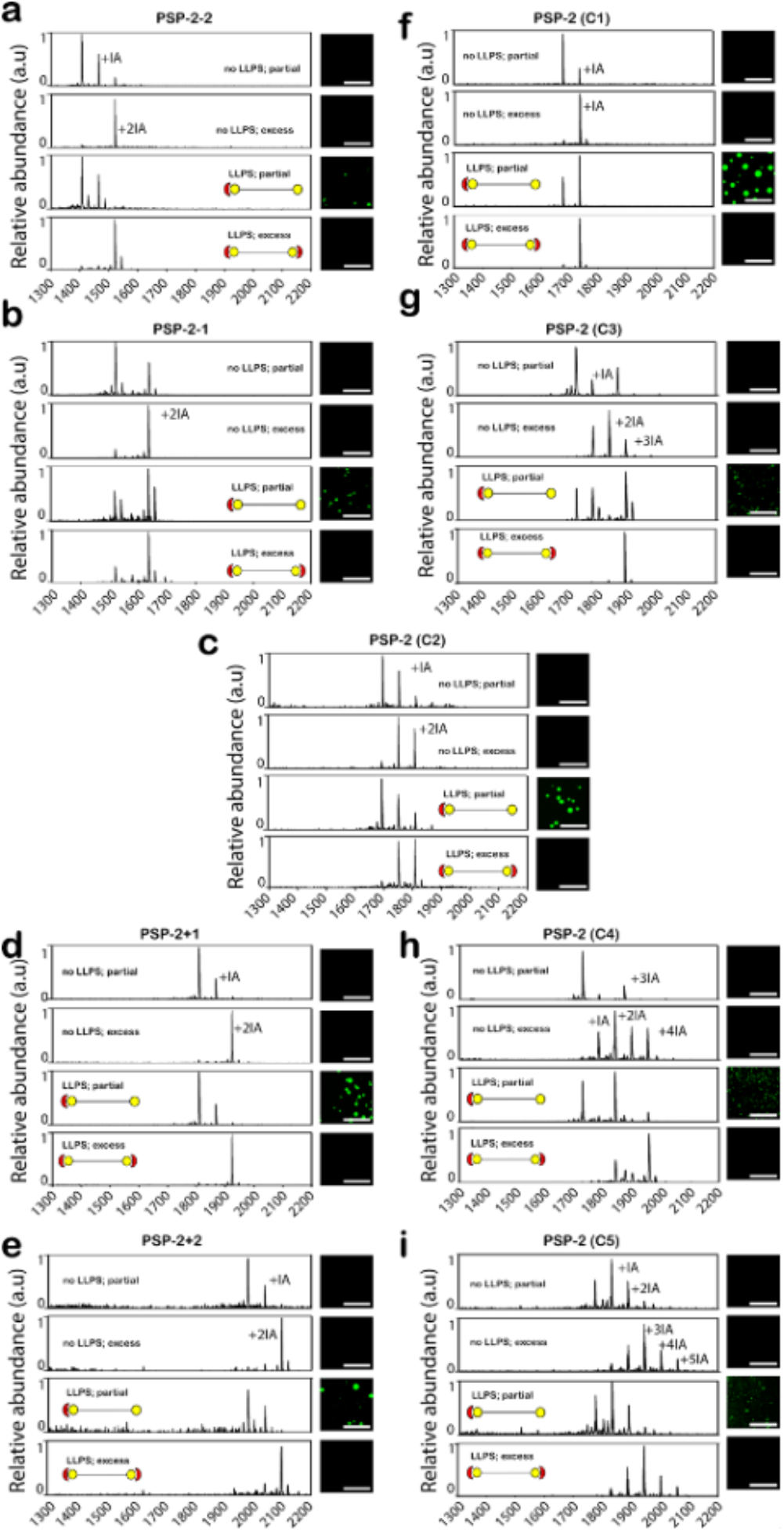
Disulfide bond obligates condensate formation. MALDI-ToF mass spectrometry spectra showing the effect of disulfide bond oxidation on condensate formation for PSP inter-cysteine length variants (a–e), and the cysteine density variants (f-i). The peptides were incubated at 5 mM concentrations for 2 h in water at room temperature (non-phase separation conditions; 20mM Tris, 50 mM NaCl, pH 8.0) either with a 0.5-fold molar excess of the capping agent, iodoacetamide (IA), for partial capping or with a 5-fold molar excess of IA for complete capping. Then, peptide stocks were diluted to a final concentration twice their respective *C_sat_* under phase-separation conditions (20 mM Tris, 2.5 M NaCl, pH 8.0).

**Figure S3.**
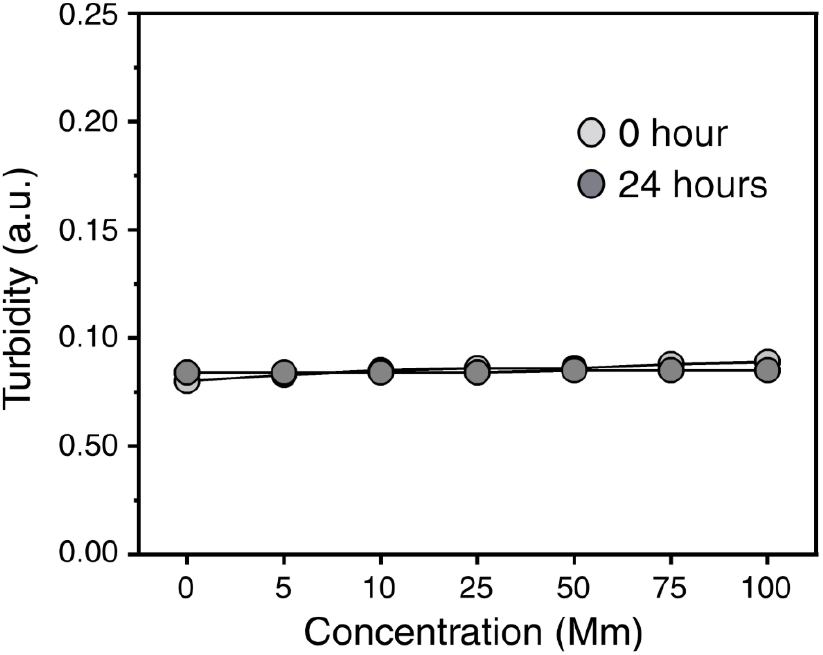
Phase diagram for the control peptide. Turbidity of the control peptide was measured (600nm) at different concentrations from 0 to 100 mM at 0 h and again after 24 h. The turbidity remained almost unchanged across the tested concentration range and between the two time points. These results suggest that the control peptide did not undergo phase separation.

**Figure S4.**
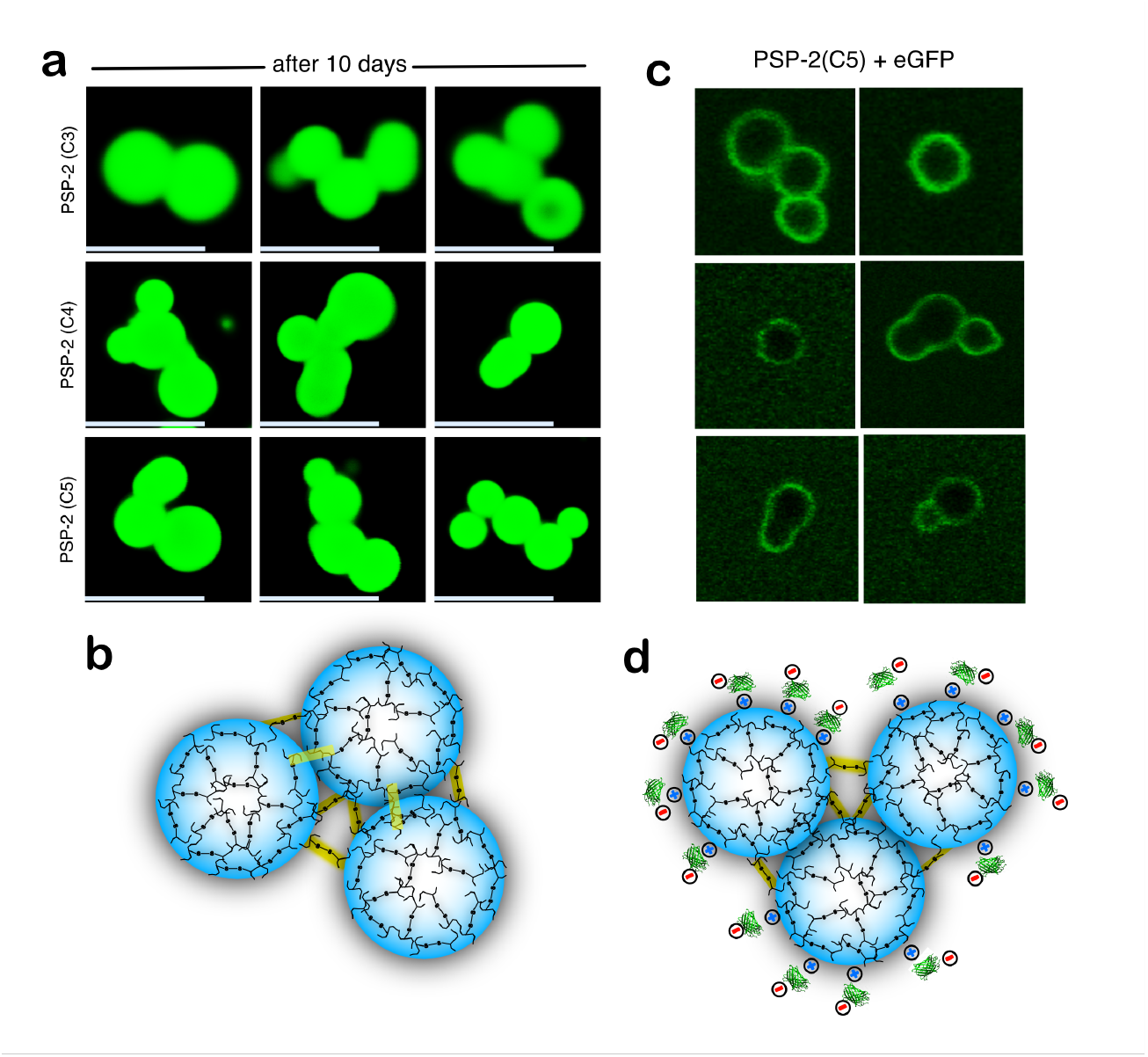
Possible mechanistic details on the behavior of high cysteine-density condensates. (a)Representative fluorescence microscopy images of PSP-2(C3), PSP-2(C4), and PSP-2(C5) condensates after 10 days, showing that the condensates remain closely associated and do not completely coalesce over time. (b) Schematic representation of a possible interaction between neighboring condensates, where increased cysteine content may promote disulfide cross-linking and stronger interactions at the condensate interface. (c) and (d) Representative fluorescence images of eGFP with PSP-2(C5) condensates showing intermolecular crosslinks via disulfide bonds.

**Figure S5.**
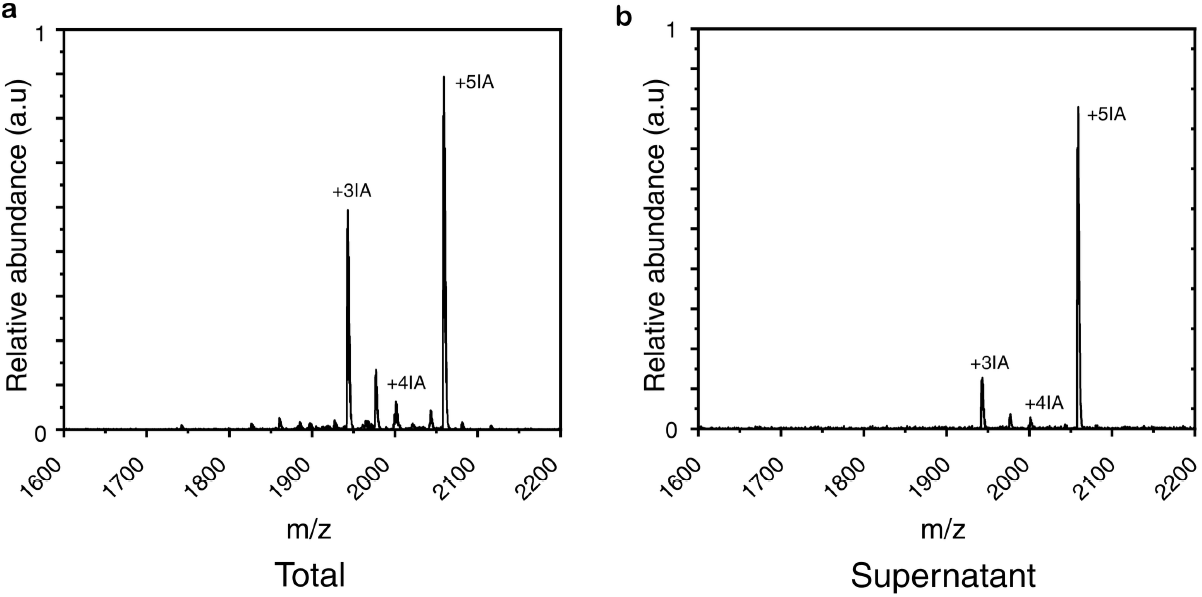
Mass spectrometric analysis of IA-capped C5 variant. Mass spectra of the IA-capped C5 peptide measured from the total sample (a) and the supernatant fraction (b). Peaks corresponding to +3IA, +4IA, and +5IA represent peptide species containing three, four, or five free cysteine residues that were capped by iodoacetamide (IA), respectively.

## Notes

### Competing Interest Statement

The authors have declared no competing interest.

